# Factors influencing the measurement of assimilation and stomatal conductance with the LI-COR 6400XT gas exchange system

**DOI:** 10.1101/494120

**Authors:** Daniel R. LeCain, Sean M. Gleason

**Affiliations:** USDA-ARS Rangeland Resources and Systems Research Unit; Fort Collins, Colorado, USA; USDA-ARS Water Management and Systems Research Unit; Fort Collins, Colorado, USA

## Abstract

Although CO_2_ and H_2_O exchange rates are often measured in experiments as indicators of physiological plant responses these “gas exchange” measurements are prone to large experimental error. Gas exchange equipment and technology have improved greatly over the past two decades which supports scrutinizing current issues of experimental error in measuring plant photosynthesis and stomatal conductance. This report shows results of a greenhouse experiment with the goal of identifying lessor understood sources of experimental error and variation in measurements with the LI-COR 6400XT gas exchange system. A variety of plant types were used to encompass differing species variation. We found significant sources of experimental error in 1) the time for initial adjustment when placing a leaf in the leaf chamber 2) the time-of-day when measuring 3) leaf age 4) having the chamber window full vs. partially full with leaf tissue 5) using a leaf chamber environment that greatly diverges from the whole plant environment 6) differing degree of experimental error depending upon plant species. A situation with multiple contributors to error would result in useless gas-exchange data. Recommendations for minimizing these experimental errors are given.

## Introduction

Although CO_2_ and H_2_O exchange rates are often measured in experiments as indicators of physiological plant responses these “gas exchange” measurements are prone to large experimental error. Certainly many actual factors influence photosynthesis and stomatal conductance, such as plant water and nutrient status, but there are additional methodological factors that contribute to error. To some degree each researcher must identify and minimize experimental error in their specific project, leading to an extensive “learning curve” for gas exchange measurements. Currently, there is a shortage of practical information to help users reduce measurement variability. A literature search shows that reports on factors influencing leaf gas exchange are somewhat dated (Long et al. 1996; Long & Bernacchi 2003). Manufacturers of gas exchange systems, such as LI-COR Biosciences (Lincoln, Nebraska, USA) typically provide instruction and documents that assist the researcher in gathering reliable gas exchange data. Although gas exchange equipment and technology have improved greatly over the past two decades there continue to be multiple sources of experimental error and uncertainty in gas exchange data. Prudent researchers typically utilize these well-understood factors:

1. Measure recently fully expanded leaves to reduce variability due to leaf age.
2. Take measurements during the peak hours of plant activity.
3. Set the gas-exchange leaf chamber conditions near “ambient”; temperature, CO_2_, humidity etc.
4. Use a non-limiting light intensity.

Improvements in the technology of leaf gas exchange equipment support taking another look at these issues. This report shows results of a greenhouse experiment with the goal of identifying lessor understood sources of experimental error and variation in measurements with the LI-COR 6400XT gas exchange system. A variety of plant species were used to include much of the existing variation in leaf size, photosynthetic systems (C_3_ & C_4_), and phylogeny. Recommendations toward balancing error with the practicality of experiments are included. The critical parameters of net CO_2_ assimilation (hereafter “assimilation”; A) and stomatal conductance (Gs) were measured. Leaf level assimilation is a fairly straight-forward measurement of CO_2_ flux (corrected for water vapor dilution), whereas stomatal conductance is a calculated parameter, requiring measurement of leaf transpiration rate, leaf temperature, and atmospheric conditions (temperature, water vapor, pressure) (Ball, Woodrow & Berry 1987). We quantified the change in assimilation and stomatal conductance resulting from various manipulations of plant and leaf chamber conditions. Following these manipulations, we assumed leaf-level equilibration when A and Gs stabilized to the chamber conditions. This was ascertained by plotting A and Gs against time using the 6400XT software-.

The experiment attempted to answer the following questions:

1. How long does it take to achieve equilibration after changing leaf chamber conditions to each of light, CO_2_, humidity and temperature?
2. Does time to equilibration vary greatly in well-watered vs. water-stressed plants?
3. How much variability is typical with changing time of day?
4. To what degree does leaf age affect gas-exchange?
5. Is there an effect of having the leaf chamber only partially filled, as is often the case when measuring plants with small leaves (even when measurements are normalized by leaf area)?
6. How independent is leaf-level functioning (within the leaf chamber) from whole plant level functioning?
7. Do these factors vary between monocots and dicots or between C_3_ and C_4_ species?

## Methods

We utilized the LI-COR Biosciences 6400XT steady-state gas-exchange system (2010 model) with the narrow leaf chamber and red/blue light source (6400-02B). Calculations were performed in the 6400XT software (OPEN version 6.2.5). When leaves were too small to fill the chamber we carefully measured leaf area and used the LI-COR recalculation spreadsheet.

Four species were grown from seed: Maize (*Zea mays* - “Flint corn”) a wide-leaved C4-type photosynthesis monocot; *Pascopyrum smithii* (Western wheatgrass) a narrow-leaved C3 monocot; Pea (*Pisum sativum* - “Lastons #9) a small-leaved C3 dicot and *Abutilon theophrasti* (Velvetleaf) a large-leaved C3 dicot. Three replications of each were sown in a commercial potting soil and grown in a greenhouse under a 12-hour photoperiod (700 to 1900 hours) with 28/20 °C day/night, 800 μmol m^−2^ s^−1^ photosynthetic active radiation (PAR), 25% relative humidity (RH) and 400 ppm [CO_2_]. Day-length was increased during this winter experiment using high pressure sodium and metal halide lamps. Plants were irrigated daily and received weekly fertilizer supplements (Dyna-gro All Pro 7-7-7; Richmond, CA). Seeds were sown on October 24, 2017 and measurements began November 24, 2017, at which time most plants were of suitable size for measurements (except Western wheatgrass). LI-COR 6400 XT measurements were performed on the same greenhouse bench as the plants were growing. Standard 6400XT startup and calibration protocol was carefully followed. Frequent “matching” of the reference and sample analysis cells was performed, especially when the CO_2_ concentration was altered. The leaf thermocouple was touching the leaf for all measurements, and therefore, the energy balance method for leaf temperature was not used. Leaf chamber CO_2_ concentration and RH were controlled with the “reference” method (incoming conditions rather than outgoing). Leaf temperature was controlled by the block method, such that the temperature of the aluminum chamber surrounding the leaf (i.e., block) was adjusted to keep leaf temperature confined to a narrow range. Generally, measurements needed to be taken within a short time window during each day. Considering that equilibration times were often significant, this usually allowed for only one measurement per species on each day.

### Trial one

How long does it take to achieve equilibration after changing leaf chamber conditions to each of light, CO_2_, humidity and temperature? Unfortunately, Western wheatgrass plants were quite small when other species were ready to measure and therefore were not included in trial one. All plants were well watered. Measurements were performed from about 1000 to 1400 hours, bracketing mid-day (preliminary tests showed that Maize was slow to reach daily peak gas exchange rates and therefore Maize was always measured last). Mid-sections of fully mature leaves of each species were clamped in the chamber at starting conditions similar to the greenhouse: 1000 μmol m^−2^ s^−1^ PAR, chamber block temperature of 24 °C, 450 ppm CO_2_ and RH of ca 25%. The chamber window was full (6 cm^2^ leaf area) for all three species. Once A and Gs had equilibrated to the chamber starting conditions the conditions were changed and monitored for a new equilibrium in response to 1) doubling the PAR, 2) doubling the [CO_2_], 3) increasing the block temperature by 8 °C, and 4) doubling the RH. We also tested the response to changing the chamber flow rate. Although flow rate is generally not considered a critical parameter, we desired to know if leaf physiology was altered at differing flow rates. Data was recorded every 3 minutes and A and Gs were plotted to determine when equilibrium to chamber conditions had been reached. Only one replication of each species was measured for trial one.

### Results

Maize was slow to equilibrate to the starting conditions, taking about 12 minutes (Figure 1A & B). The 6400XT achieved adjustments in chamber environment very quickly for all parameters, and therefore was not a delaying factor in the analyses. The Maize response to increasing PAR and increasing CO_2_ were very fast (less than 3 minutes) (Figure 1 A & B). The Gs response to doubling CO_2_ was marked and quickly detected (Figure 1B). As predicted there was little response to increased flow rate. Assimilation slowly increased with increasing leaf chamber temperature, taking about 12 minutes to resume equilibration. Surprisingly, Gs had little response to higher temperature, and therefore higher vapor pressure deficit (VPD). Similarly, there was little response to increased RH, and therefore lower VPD (Figure 1A & B).

**Figure 1.**
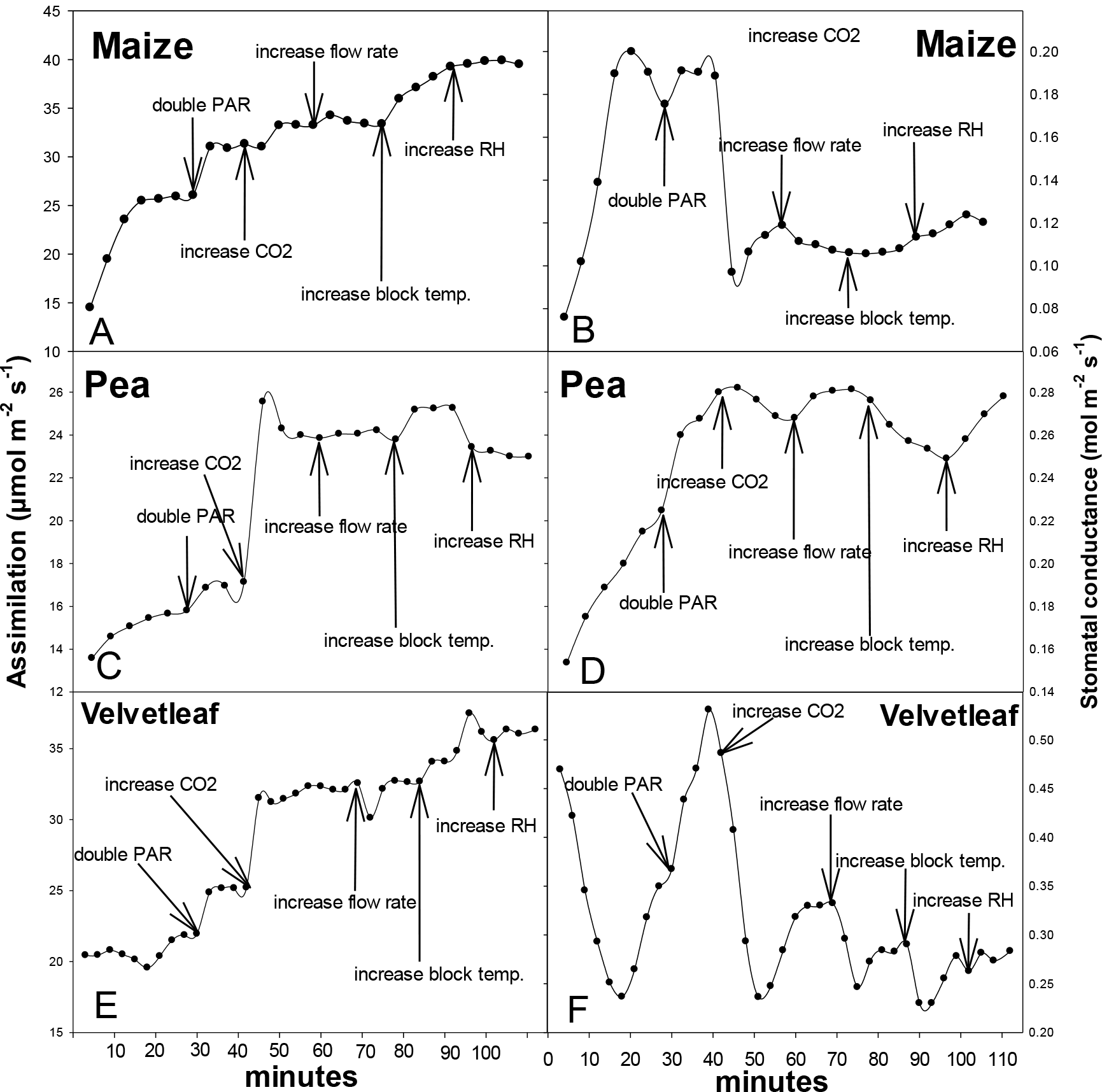
The response of Maize, Pea and Velvetleaf assimilation (A, C, E) and stomatal conductance (B, D, F) to changes in the leaf chamber environment (photosynthetically active radiation (PAR), CO_2_ concentration, flow rate, chamber temperature and relative humidity (RH) using the LiCor 6400XT gas exchange system.

Pea leaves were also slow to equilibrate to starting conditions, taking about 15 minutes (Figure 1C & D). Similar to Maize, equilibration to changes in chamber environment were quite fast; about 3 minutes for an increase in PAR, temperature, RH and [CO_2_]. There was a small change in Gs in response to higher flow rate, but not A (Figure 1 C & D). The stomatal conductance response to warmer and higher RH chamber conditions was slower than the assimilation response. In general assimilation equilibrated more quickly than stomatal conductance.

Velvetleaf was very slow to equilibrate to starting conditions, particularly in stomatal conductance, taking about 21 minutes to reach a steady-state (Figure 1E & F). Assimilation responses to chamber changes were fast but Gs responses were slow and erratic. In particular, Velvetleaf Gs reacted to a change in flow rate that was not fully compensated for in the calculations.

Despite a chamber environment very similar to the growth environment, all three species required a long time to reach full equilibrium to the leaf chamber (ca 16 minutes). It is typical to report allowing 5 minutes or less for initial equilibration to a gas-exchange chamber (LICOR 6400XT manual). Our results suggest up to a 25% error if data are logged at 5 minutes. Our test shows that subsequent changes to chamber conditions are fast, re-equilibrating in about 3 minutes. Furthermore, Gs generally equilibrated more slowly than A, particularly for VelvetLeaf, where Gs exhibited a slow and erratic equilibration. Stomatal changes to vapor pressure may be particularly slow (Turner et al. 1984). VelvetLeaf also seemed to be affected by a change in flow rate, which in theory is completed adjusted for in the calculations.

### Trial two

Does equilibration time vary greatly in well-watered vs. water-stressed plants? Trial two was performed at the end of the study period but is presented here for comparison with trial one. Pots were allowed to dry to about 20% of their saturated weight. This seems extreme but the potting soil used had a very large water holding capacity. This degree of soil drying was just above the visible wilting point for all species. In order to shorten the measurement duration, only chamber PAR and [CO_2_] were altered. Surprisingly, Maize A and Gs rates were nearly the same as in well-watered plants, suggesting that we did not achieve the desired water stress, and therefore we do not report data for these species here.

### Results

Pea leaves of water-stressed plants responded to chamber changes in PAR and [CO_2_] in parallel to well-watered plants, albeit with lower A and Gs rates (Figure 2A & C). As with well-watered plants, stressed Pea plants were quite slow to equilibrate to starting conditions and were still adjusting slightly after 18 minutes. There was a slightly different response in Gs to increasing [CO_2_] in water stressed plant, with a slow decrease in Gs which was not seen in well-watered plants. However this is an expected response to higher [CO_2_] in most plants (Figure 2C).

**Figure 2.**
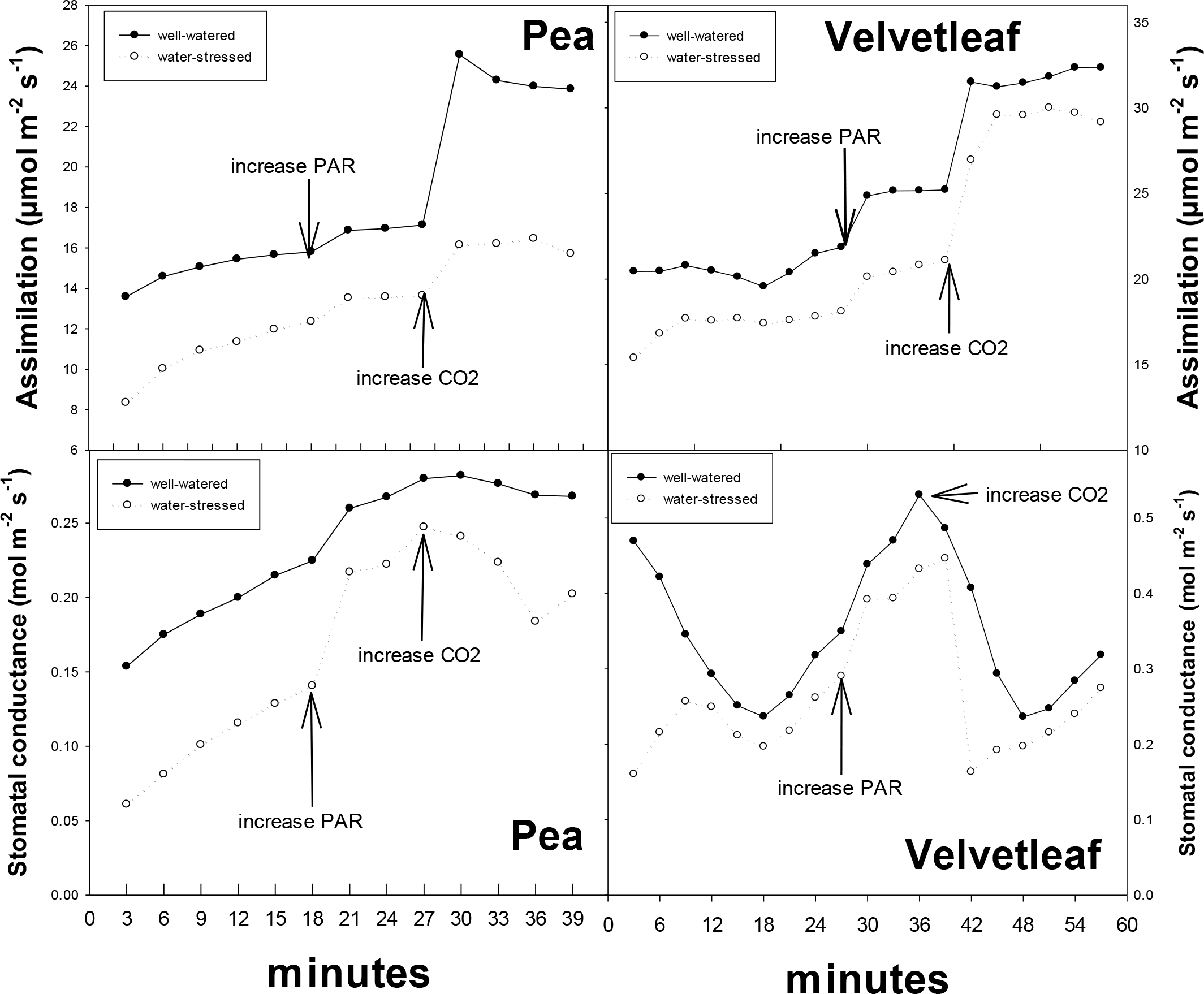
The response of Pea and Velvetleaf assimilation and stomatal conductance to changes in photosynthetically active radiation (PAR) and CO_2_ concentration in well-watered vs. water-stressed plants using the LiCor 6400XT gas exchange system.

Water-stressed Velvetleaf responded similarly to changing chamber conditions as the well-watered plants (Figure 2B & D). Again, it required a long time for Gs to equilibrate to starting conditions (ca 20 minutes), however the initial slope of the adjustment was opposite in water-stressed plants (Figure 2D).

Although we have a limited dataset, it appears that plants subjected to mild water-stress will equilibrate to changes in leaf chamber environment in a similar way as well-watered plants. The initial equilibration to the leaf chamber will be quite slow, but subsequent changes in environment will be compatible with a 5 minute logging interval. Different species and degrees of water stress could have contrary results to ours.

### Trial 3

How much variability is typical with time of day? Leaves of all four species were measured early in the photoperiod and the leaf area that was in the chamber was marked and returned to the chamber at six different times during the photoperiod. Therefore the exact section of the leaf was measured over the course of a day. Plants were kept well-watered throughout. Chamber conditions were similar to the greenhouse environment except that saturating light was used: 2000 μmol m^−2^ s^−1^ PAR, 450 ppm [CO_2_], 25 °C and 25% RH during all measurements.

### Results

Assimilation decreased late in the photoperiod for all species but there were differences among species (Figure 3). Maize A did not decrease until about 10 hours into the photoperiod (Figure 3A). Maize also did not achieve peak A until after 0900 hour. Pea assimilation decreased shortly after midday (Figure 3B). Velvetleaf declined slightly over the course of the day (Figure 3C). Western Wheatgrass A did not decline until about 10 hours into the photoperiod (Figure 3D).

**Figure 3.**
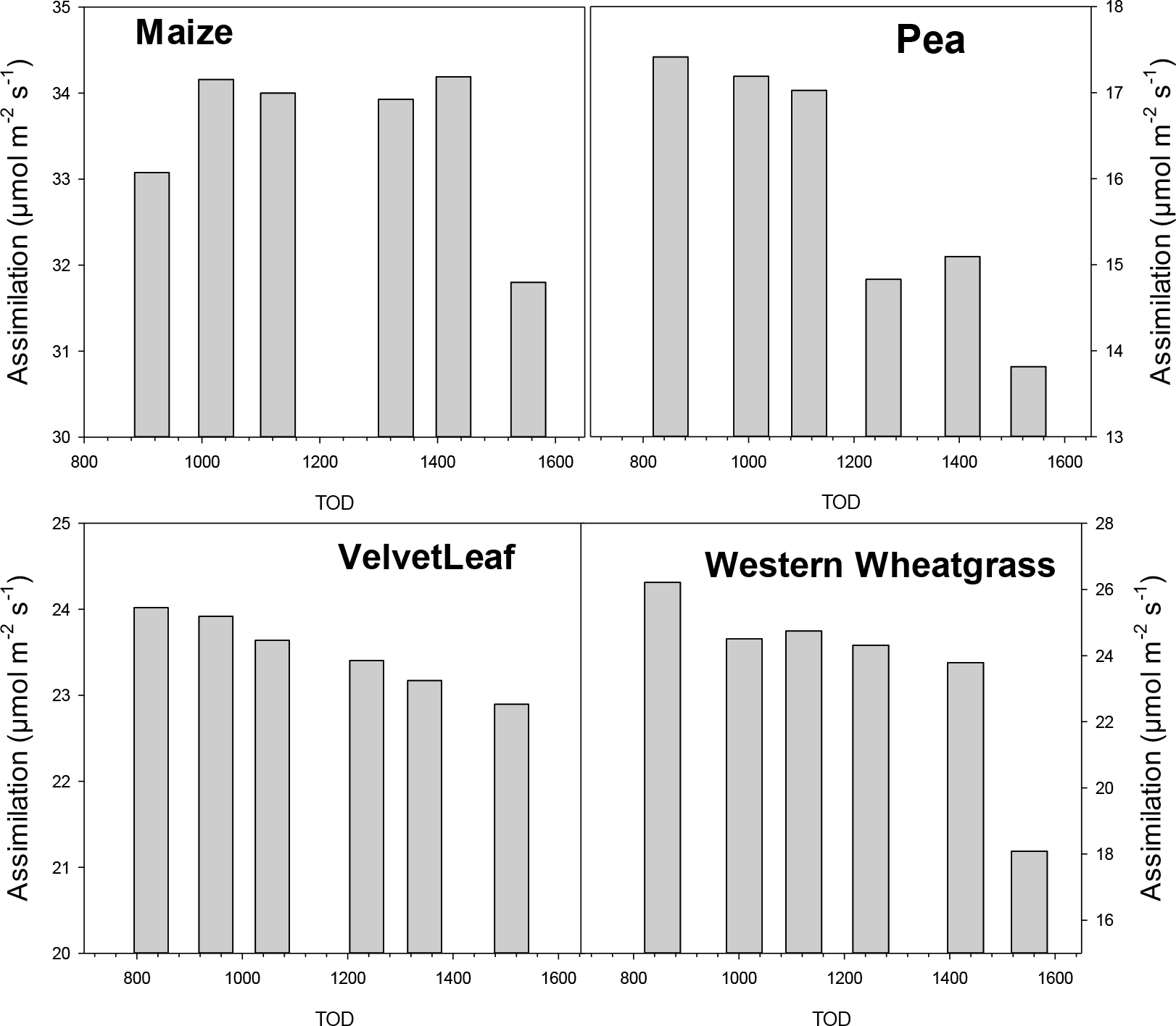
The influence of time-of-day of on the assimilation rate of Maize, Pea, Velvetleaf and Western wheatgrass using the LiCor 6400XT gas exchange system.

Stomatal conductance rates over the photoperiod were less consistent than assimilation (Figure 4). Pea Gs behaved similarly to A with a strong decline about 10 hours into the photoperiod (Figure 4B), however other species had more erratic Gs rates. The striking finding was that Gs of Maize was lower in the morning vs after midday, although only by ca 15% (Figure 4A; note the small range on the Y axis). In general, consistent measurement of Gs was more difficult than A.

**Figure 4.**
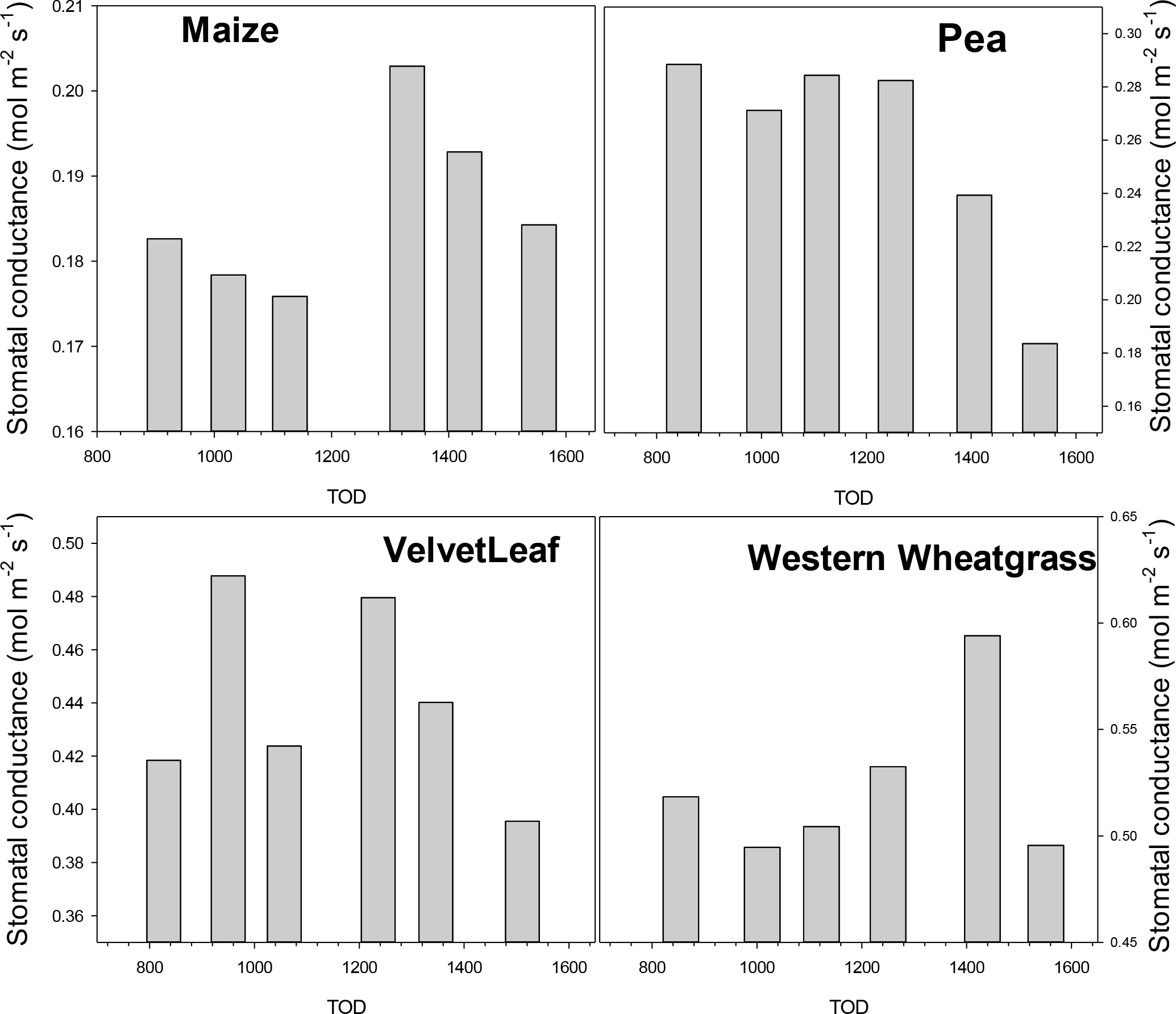
The influence of time-of-day on the stomatal conductance of Maize, Pea, Velvetleaf and Western wheatgrass using the LiCor 6400XT gas exchange system.

### Trial 4

To what degree does leaf age affect gas-exchange? For this question leaves were labelled with the date when they became fully expanded (as plant growth progressed). When there was a 20 day range in leaf age these leaves were measured under chamber conditions similar to the greenhouse environment as in Trial 1. Western wheatgrass was not measured due to difficulty in determining leaf age. We report only assimilation data (Figure 5), but note that stomatal conductance responded similarly in all cases.

**Figure 5.**
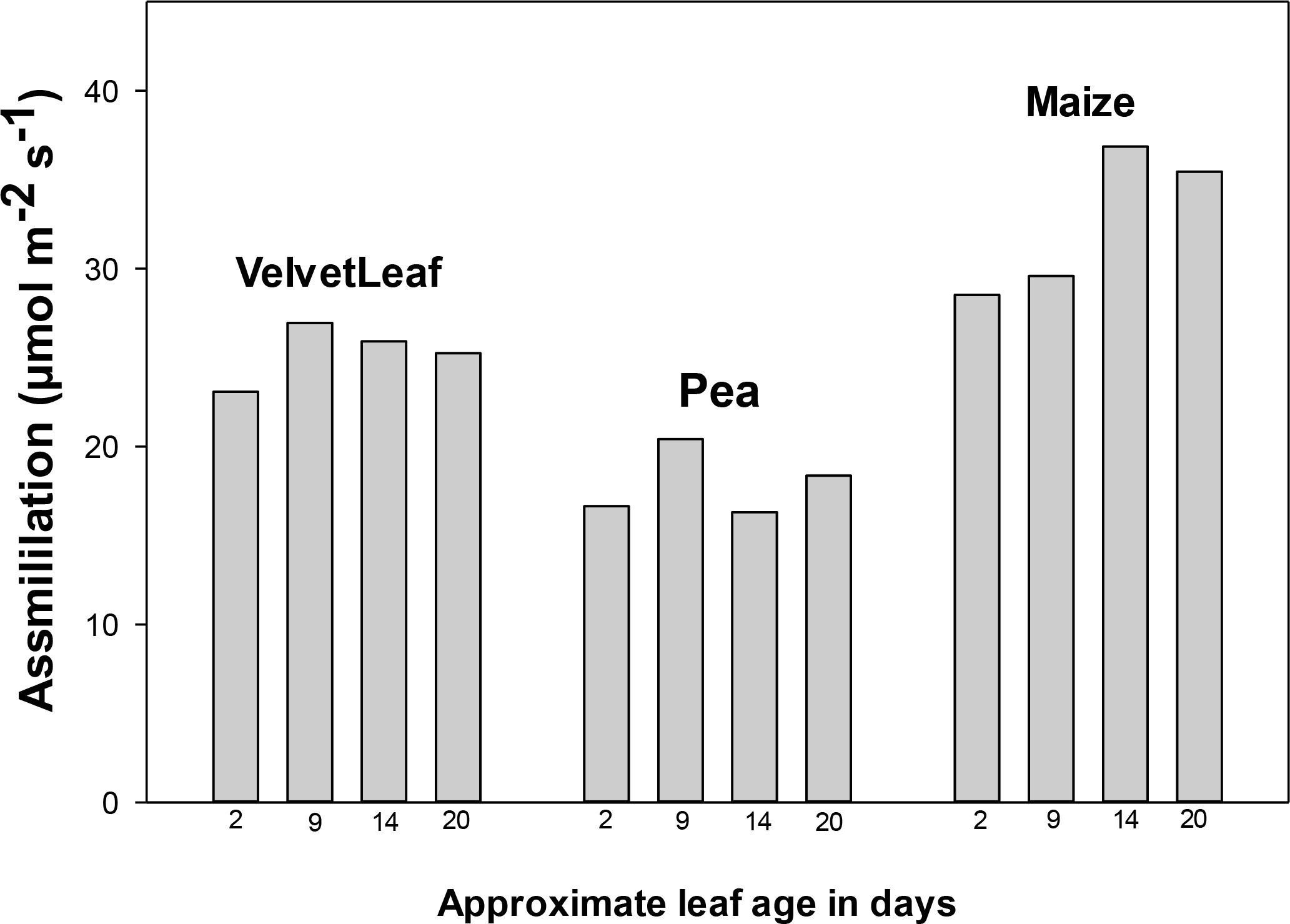
The influence of leaf age on the assimilation rate of Velvetleaf, Pea and Maize using the LiCor 6400XT gas exchange system.

### Results

Under greenhouse conditions with plentiful irrigation and nutrients, leaf age, ranging by about 18 days, had minor effects on assimilation rates (Figure 5). However it is noteworthy that Maize A did not peak until leaves were over 9 days old. This is somewhat contrary to the perception that gas-exchange rates are highest in young and recently expanded leaves (Long et al. 1996). However this again may be a response to the general low light greenhouse conditions during the winter and C_4_ photosynthesis of Maize.

### Trial 5

Is there an effect of having the leaf chamber window only partially filled (even with measurement of leaf area)? Many wild and cultivated species exhibit leaves that are smaller than the chamber window of most gas exchange systems. This situation results in only a partially filled chamber, and subsequently, requires the recalculation of both A and Gs. However, it remains uncertain if the air flow through a partially filled chamber (relative to a full chamber) differs, or results in meaningfully different measurements of A or Gs. At the very least researchers are often encouraged to standardize the amount of leaf in a partially filled chamber window. Trial 5 was a simple test of this issue using Maize and Pea leaves. This trial was performed on two replicates for each of Maize and Pea.

For Maize, a large leaf was chosen that filled the entire chamber (6 cm^2^). This leaf was allowed to fully equilibrate, and A and Gs were recorded. Then, a slit was made along the length of the leaf to create a more narrow section of leaf, which was then quickly returned to the chamber. The chamber area filled by the cut Maize leaf was determined from leaf width (measured with digital calipers) and chamber length.

For Pea, a large leaf that filled the chamber window was measured at full equilibrium, then a smaller leaf that was nearby on the same stem was measured. The smaller leaf area was measured on the LiCor area meter (3000A).

### Results

Pea leaves that only partially filled the chamber window had slightly lower A and Gs (Figure 6). We note that these differences could have resulted from random variation across leaves, although the large and small leaves were very close on the stem. Maize leaves that partially filled the chamber had slightly higher A and Gs rates, and this was measured on the very same leaf tissue. It is possible that the cut edge of the Maize leaf could have influenced this test, however the cut surface area was very small compared to the intact leaf surface area. It is also possible that air circulation around both sides of the leaf was slightly different in the partially filled chamber and that this had a small effect on the measurements.

**Figure 6.**
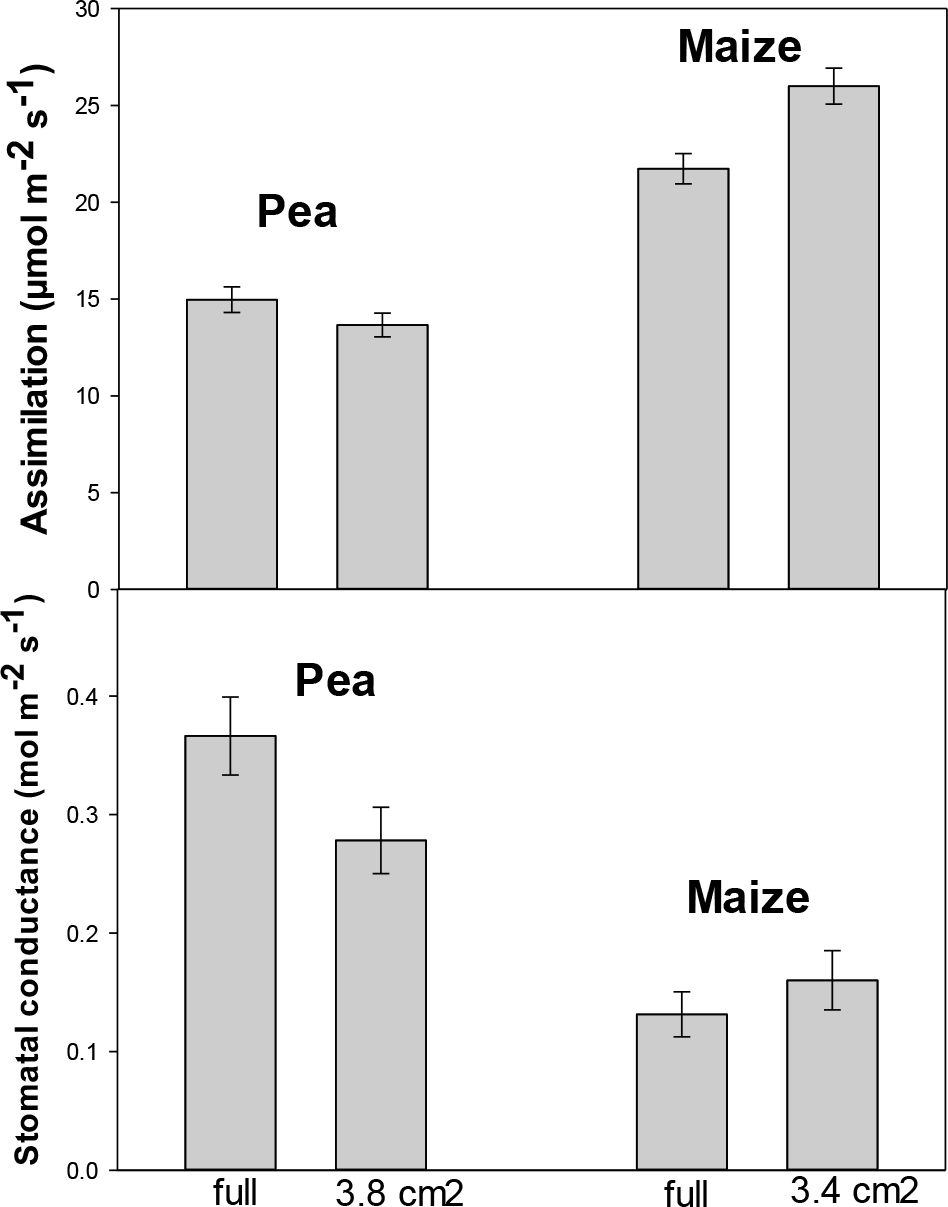
The influence of a full vs. partially full leaf chamber window on assimilation and stomatal conductance in Pea and Maize using the LiCor 6400XT gas exchange system.

### Trial 6

How independent is leaf-level functioning (within the leaf chamber) from whole plant level functioning? To have confidence in gas exchange measurements the leaf portion that is in the leaf chamber must be responding primarily to the chamber environment. Chamber conditions that diverge from the conditions of the whole plant, such as measurement at elevated CO_2_, will hopefully have the desired influence on the leaf section in the chamber, rather than the conditions the remainder of the plant is exposed to (outside the leaf chamber). One must also assume that the chamber is not leaky to the external environment (Rodeghiero et al. 2007). Trial 6 tested the independence of the leaf portion in the chamber and the whole plant. Whole plants were placed in controlled environment growth chambers (Conviron BDR16, Winnipeg, Manitoba, Canada) and the growth chamber conditions were altered under steady 6400XT leaf chamber conditions. The starting growth chamber conditions were 25C, 30% RH, 400 μmol m^−2^ s^−1^ PAR and 420ppm CO_2_. These starting conditions were changed to 35 °C, 80% RH, 800 μmol m^−2^ s^−1^ PAR, zero PAR (lights off), and 750 ppm CO_2_. Note that these manipulations were not always in the same order for each species, although we do not expect this affect the responses. 6400XT leaf chamber conditions were 25C, 30% RH, 1000 PAR and 420 CO_2_ and were unaltered. Because the growth chamber had to be closed during these tests, the 6400XT console was viewed through a window in the growth chamber door and when equilibrium was reached the data were quickly recorded, the growth chamber door was closed and the next growth chamber manipulation began. Growth chamber changes in PAR, [CO_2_] and RH were quite fast, however it took about 15 minutes for the growth chamber to increase by 10 C.

### Results

Maize single leaf gas exchange was strongly influenced by a 10 degree increase in the temperature of the whole plant; both A and Gs increased even though the leaf chamber temperature remained at 25C (Figure 7A). Maize also exhibited a small but temporary response to both increasing and decreasing the light on the whole plant. There was no effect of whole plant CO_2_ environment, nor to increasing the relative humidity.

**Figure 7.**
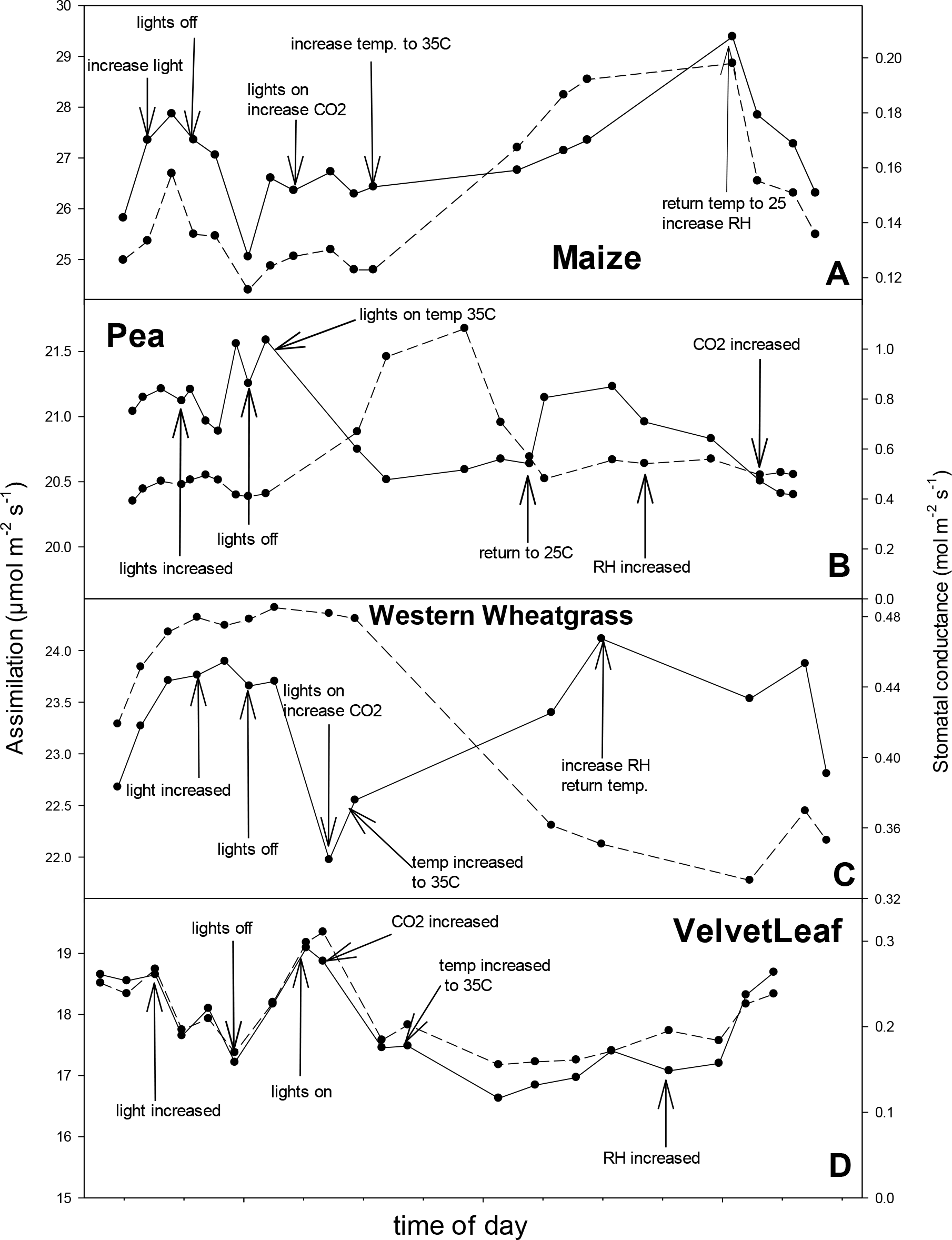
The influence of changes in the whole plant environment (growth chamber light level, CO_2_ concentration, temperature and relative humidity) on assimilation (solid line) and stomatal conductance (dashed line) while holding the leaf chamber environment steady in Maize, Pea, Western wheatgrass and Velvetleaf using the LiCor 6400XT gas exchange system.

Pea leaves had an especially strong and positive Gs response to an increase in whole plant temperature (note Y axis range), however this response was temporary (Figure 7B). Pea exhibited a small but noticeable decrease in assimilation in response to increased whole-plant temperature. Similar to Maize, there appeared to be very small and temporary responses to changing light, [CO_2_] and RH.

Western Wheatgrass exhibited an increase in assimilation, but a decrease in stomatal conductance in response to increasing whole plant temperature (Figure 7C). There was little effect of changing whole plant PAR, CO_2_ nor RH.

Assimilation in VelvetLeaf declined slightly in response to higher [CO_2_] and higher whole plant temperature, however, there was much variation in these measurements (Figure 7D). An increase in whole plant RH elicited a rather large increase in A and Gs. Assimilation and stomatal conductance tracked well in Velvetleaf.

Single leaf gas-exchange was affected by divergence in whole plant environment in all four species. In particular, leaves held at constant climate conditions within the 6400XT chamber exhibited meaningful changes in functioning when whole-plant temperature was increased by 10 °C. Maize and Western wheatgrass assimilation exhibited a positive response to increasing whole plant temperature, whereas the two dicotyledons exhibited negative responses. In general, divergence in whole plant vs leaf section functioning were very small for changes in [CO_2_], PAR and RH. However, VelvetLeaf and Western Wheatgrass exhibited small negative responses to increasing whole plant [CO_2_].

Finally, we ask, do measured responses differ between monocots and dicots, or between C_3_ and C_4_ plants? Certainly, what we can interpret from this small set of species is quite limited. Of note however, is that Maize stomatal conductance was very low during the morning hours. This could be due to the relatively low light environment in the greenhouse in the winter, since C_4_ plants perform well under conditions of high heat and irradiance. There was also some indication that VelvetLeaf responded more erratically than the other species when leaf chamber conditions where changed. In particular, Gs exhibited much variation in this species (see Figure 3 & 7). We suspect that this result may have arisen due to the very thin leaves of this species. VelvetLeaf also responded to increasing flow rate, i.e., the 6400XT did not appear to fully normalize A and Gs for flow rate. This suggests that boundary layers may have been dissimilar between Velvetleaf and the other species.

Not surprisingly, results varied with species, emphasizing that researchers should perform a thorough trial run of gas exchange measurements on their species of interest. Field measurements will be very difficult to standardize for leaf age, full vs. partially full chamber, whole plant vs. chamber conditions, plant water and nutrient status and time of day influences, all adding to the high variability in gas exchange data.

## Recommendations

Table 1 summarizes the potential error in assimilation for each of the investigated factors. There was some evidence that the experimental error may have been higher in stomatal conductance measurements. Certainly, a worse-case scenario involving multiple factors could easily result in useless gas-exchange data. In addition to these method concerns there are significant sources of error in actual experiments due to plant phenology, variable soil water and nutrition, intra- and interspecific genetic variation etc. Additionally, unique stem and leaf morphologies may add another layer of difficulty, resulting in difficult clamping and sealing of leaves (Rodeghiero et al. 2007). Researchers must always attempt to balance known and unknown experimental error with the practical aspects of gathering gas-exchange data. Limited personnel, time and even weather can often dominate protocol decisions. We recommend that researchers conduct as thorough preliminary tests as possible prior to gathering important gas-exchange data. At the very least, standardization of gas-exchange protocols is important for: 1) time of day, 2) temperature, 3) leaf age, 4) leaf size in the chamber, 5) criteria for determining equilibrium, and 6) divergence from whole plant conditions. Thorough preliminary tests and data analysis will likely reduce the amount of undesired variation in gas-exchange measurements. Increasing the number of replications per study group should add additional confidence to gas-exchange data.

**Table 1.**
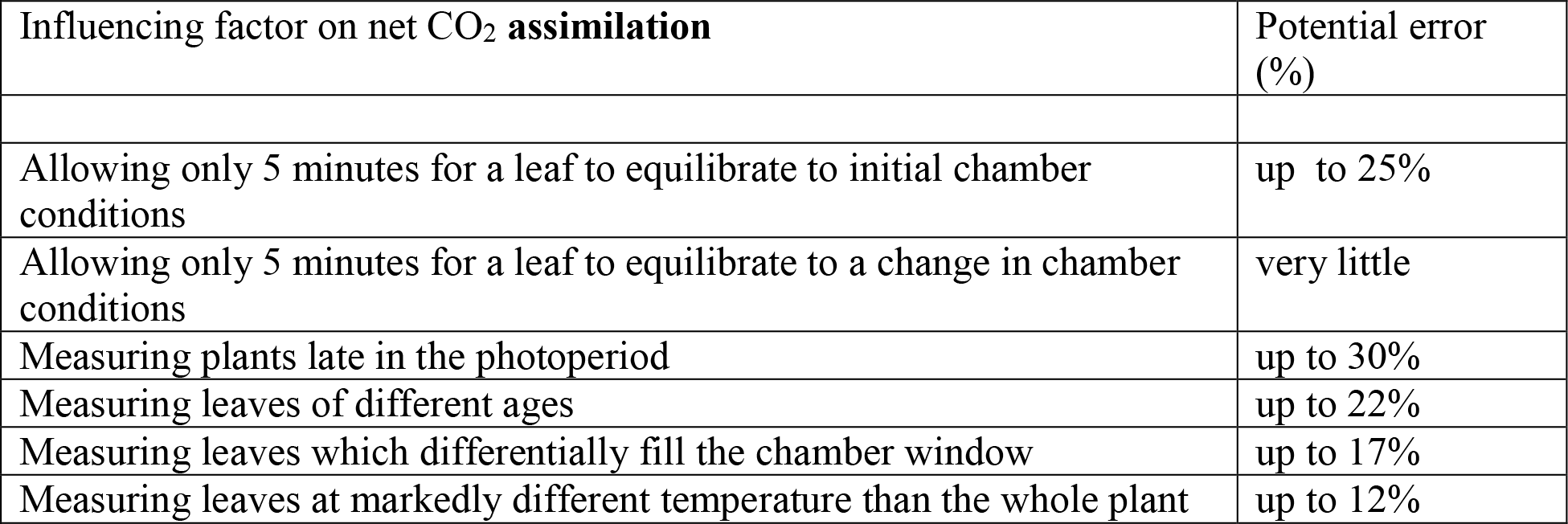

